# Differential translation of mRNA isoforms underlies oncogenic activation of cell cycle kinase Aurora A

**DOI:** 10.1101/2023.03.13.532331

**Authors:** Roberta Cacioppo, H. Begum Akman, Taner Tuncer, A. Elif Erson-Bensan, Catherine Lindon

**Author notes:** To whom correspondence should be addressed. Email: Roberta Cacioppo or Catherine Lindon.

## Abstract

Aurora Kinase A (AURKA) is an oncogenic kinase with major roles in mitosis, but also exerts cell cycle- and kinase-independent functions linked to cancer. Therefore control of its expression, as well as its activity, is crucial. A short and a long 3’UTR isoform exist for AURKA mRNA, resulting from alternative polyadenylation (APA). We initially observed that in Triple Negative Breast Cancer, where AURKA is typically overexpressed, the short isoform is predominant and this correlates with faster relapse times of patients. The short isoform is characterized by higher translational efficiency since translation and decay rate of the long isoform are targeted by *hsa-let-7a* tumor-suppressor miRNA. Additionally, *hsa-let-7a* regulates the cell cycle periodicity of translation of the long isoform, whereas the short isoform is translated highly and constantly throughout interphase. Finally, disrupted production of the long isoform led to an increase in proliferation and migration rates of cells. In sum, we uncovered a new mechanism dependent on the cooperation between APA and miRNA targeting likely to be a route of oncogenic activation of human AURKA.

## INTRODUCTION

Aurora Kinase A (AURKA) is a critical positive regulator of the mitotic phase of the cell cycle (1). AURKA also plays additional cancer-promoting roles in cell proliferation, survival, migration, and cancer stem cell phenotypes, some of which in interphase and in a kinase-independent manner (2). AURKA expression follows a strict cell cycle-dependent pattern, with both protein and mRNA levels extremely low in G_1_ phase, increasing in S phase, and peaking at G_2_ phase until mitosis (3). High expression of AURKA is strongly associated with cancer progression, drug resistance and poor prognosis, justifying why oncogenic AURKA represents a renowned target of anti-cancer drugs (4), and making evident that oncogenic roles of AURKA are prompted by its highly sustained levels of expression.

AURKA overexpression in human cancers is known to be caused by elevated gene copy number, enhanced transcription or increased protein stability. Dysregulation of translation is also linked to disease and contributions of dysregulated translation to cancer phenotypes are increasingly reported (5) (6). Despite some modest evidence suggests that modulation of AURKA translation is relevant in disease (7) (8), control of AURKA expression at the level of translation is widely understudied compared with control of its transcription and mRNA processing (3). For example, it is not clear whether AURKA mRNA undergoes translational activation and/or inhibition during the cell cycle, and the precise timing, extent or regulators of these processes remain unexplored.

The process of cleavage of the 3’end of precursor mRNAs (pre-mRNAs) and concomitant addition of a poly(A) tail represents one key event aiding the maturation of mRNAs, termed cleavage and polyadenylation (C/P) (9). The cleavage site is typically preceded by a polyadenylation signal (PAS), located 10–30 nucleotides (nt) upstream, and by UGUA and U-rich motifs, whereas it is typically followed by U- and GU-rich motifs. Altogether, these elements constitute the C/P site (10). Most human pre-mRNAs contain multiple C/P sites (11), enabling alternative cleavage and polyadenylation (APA) and, thus, distinct expression of transcript isoforms for the same gene. A search using PolyA_DB (12) indicates presence of two C/P sites with canonical PASs (AATAAA) on AURKA 3’ untranslated region (3’UTR) (**Figure 1A**). This fostered our hypothesis that AURKA mRNA could be subjected to tandem 3’UTR APA, resulting in two 3’UTR isoforms that differ in length. It is currently unknown which AURKA PAS is preferentially used in which cellular context or whether a 3’UTR isoform switch can be modulatable.

**Figure 1:**
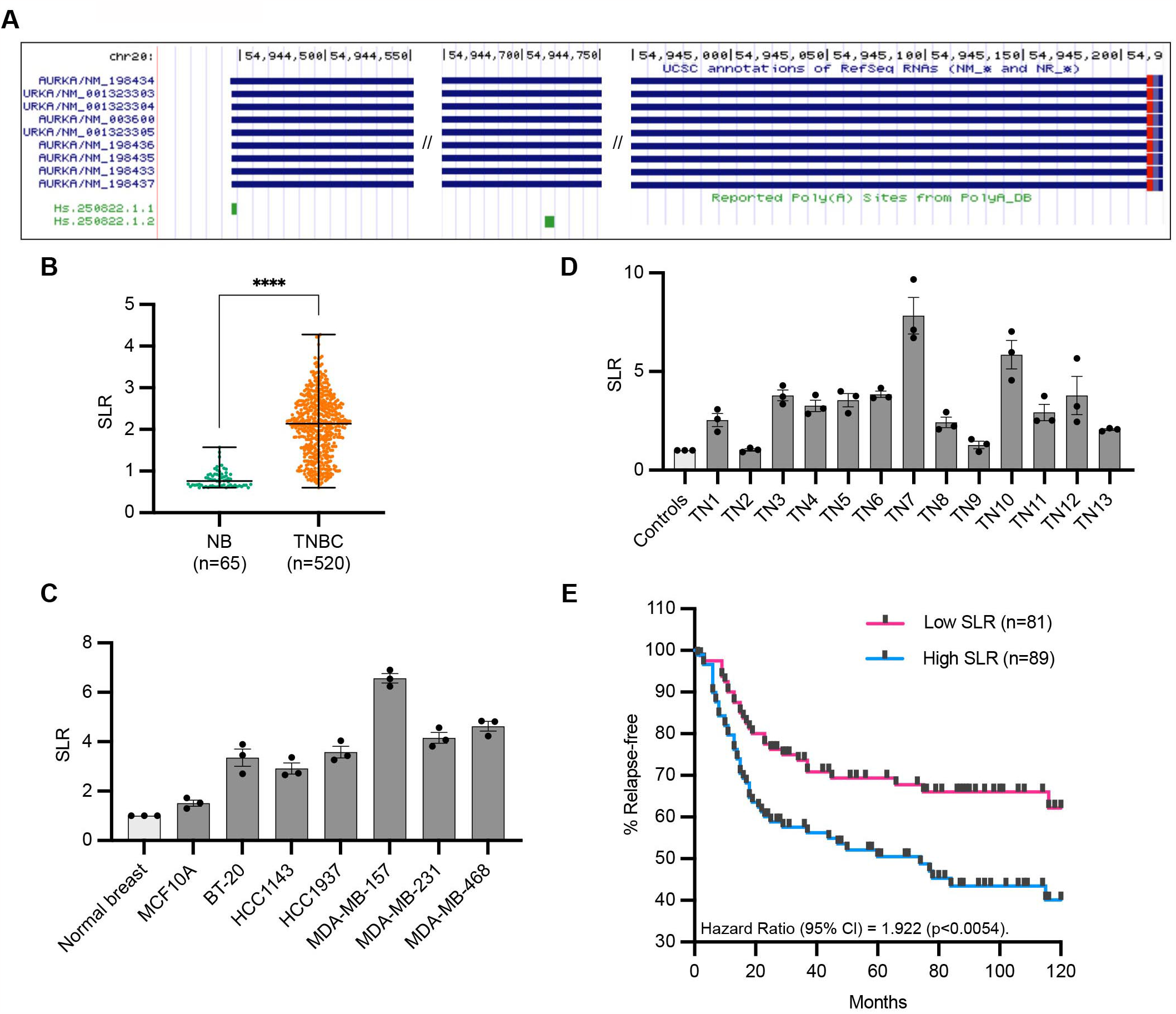
Increased short/long ratio (SLR) of AURKA APA isoforms in TNBC. **(A)** AURKA transcript isoforms (USCS Genome Browser). AURKA gene is located on (-) strand. **(B)** Median and range of SLR values for AURKA 3’UTR obtained using APADetect. Mann-Whitney test; ****, p<0.0001. **(C)**,**(D)** RT-qPCR analysis of SLR of AURKA 3’UTR in TNBC cell lines (C) and patient samples (D). SDHA used as reference gene. TN, tissue number. **(E)** Relapse-free survival rates of TNBC patients with high (highest 25%) or low (lowest 25%) AURKA SLRs. p value determined by log-rank test.

APA is involved in most cellular processes and is often altered in cancer (9). Clinically, human cancers are characterized by unique profiles of alternative 3’UTRs that can be exploited for classification of distinct cancer subtypes (13) (14), and associations between 3’UTR shortening and poor prognosis (15) or drug sensitivity (16) have been detected. At the molecular level, a strong positive association between expression of short 3’UTRs, increased protein levels and proliferative states has been frequently reported (17) (18) (19) (20) (21). Such genome-wide 3’UTR shortening sustains cancer cell behavior by removing repressor sequence elements from the 3’UTR of oncogenic mRNAs, for example microRNA (miRNA) binding sites (17) (18) (19), or alternatively by inactivating tumor suppressors through suppression of their expression (22) (23).

The role of miRNAs in regulating cell cycle genes and the relevance of this regulation in cancer are well understood (24) (25). Few miRNAs have been pointed to as regulators of AURKA mRNA but, importantly, reported cases of miRNA targeting of AURKA occur in those cancers where AURKA overexpression is a promoting factor or a marker of poor prognosis (26) (27) (28) (29). Regardless, none of these studies consider the existence of distinct AURKA 3’UTR isoforms in their experimental design of targeting assessment. The *hsa-let7* miRNA family comprises 11 closely related genes that map in chromosomal regions that are typically deleted in human tumors and, given their pathogenic downregulation in cancer, they are classified as tumor suppressors (24) (30). Roles for *hsa-let-7* in breast tumor growth and metastasis have been proposed (31) (32) and a correlation between *hsa-let-7a* expression and clinical variables has been detected in TNBC (33) (34).

AURKA was classified within the Triple Negative Breast Cancer (TNBC) subtype with the highest median index of 3’UTR shortening events (14) (35), and also undergoes 3’UTR shortening in poor-prognosis patients of breast and lung cancer (15). Importantly, AURKA overexpression in TNBC represents a marker of early recurrence, poor prognosis, and shorter overall survival (36) (37). However, the correlation between AURKA PAS usage, protein expression and pathological cell behavior has not been explored for this or other biological contexts, nor at the molecular level. In this study, we uncover a molecular mechanism leveraging the cellular ratio of APA isoforms and their different translational program during the cell cycle to control acquisition of AURKA oncogenic potential.

## RESULTS

### Increased short/long ratio of AURKA APA isoforms in TNBC

In a preliminary study using the APADetect *in silico* tool, we analysed publicly available microarray data to identify changes in AURKA 3’UTR isoform abundance in tissues (35). 520 comparable datasets for TNBC samples came from GSE31519 (48) and 65 histologically normal epithelium and cancer-free prophylactic mastectomy patients were used: 32 from GSE20437 (49), 12 from GSE9574 (50), 7 from GSE3744 (51), 6 from GSE6883 (52), 5 from GSE26910 (53), and 3 from GSE21422 (54). The analysis revealed increased short/long ratio (SLR) of AURKA 3’UTR isoforms in TNBC compared to normal breast tissues (**Figure 1B**). Higher SLR was confirmed by RT-qPCR in multiple TNBC cell lines (**Figure 1C**). Furthermore, RT-qPCR analysis of normal and TNBC patient cDNAs from Origene Breast Cancer cDNA array IV (BCRT504) also showed higher AURKA SLR in TNBC samples compared to normal (**Figure 1D**). In addition, the shortening of AURKA 3’UTR correlated with faster relapse times in TNBC patients (clinical data from (48)) (**Figure 1E**). These results therefore suggest a potential oncogenic role of AURKA APA in breast cancer worth further investigations.

### AURKA shows 3’UTR isoform-dependent protein expression

To probe AURKA APA isoform-dependent protein expression, we developed a single-cell expression sensor suitable for experiments in live cell. The construct independently expresses Venus and mCherry fluorescent proteins via a constitutive bi-directional promoter (**Figure 2A**). The coding sequence (CDS) of Venus is flanked by AURKA UTRs, whereas that of mCherry lacks regulatory regions and is therefore used to normalize for transfection efficiency. To test for APA-sensitive expression, we alternatively mutated the distal (d) or proximal (p) PAS on the reporter 3’UTR, to generate different 3’UTR isoforms (SHORT and LONG, respectively). Constructs lacking AURKA UTRs (Δ) and expressing AURKA wild-type UTRs (WT) were used as controls.

**Figure 2:**
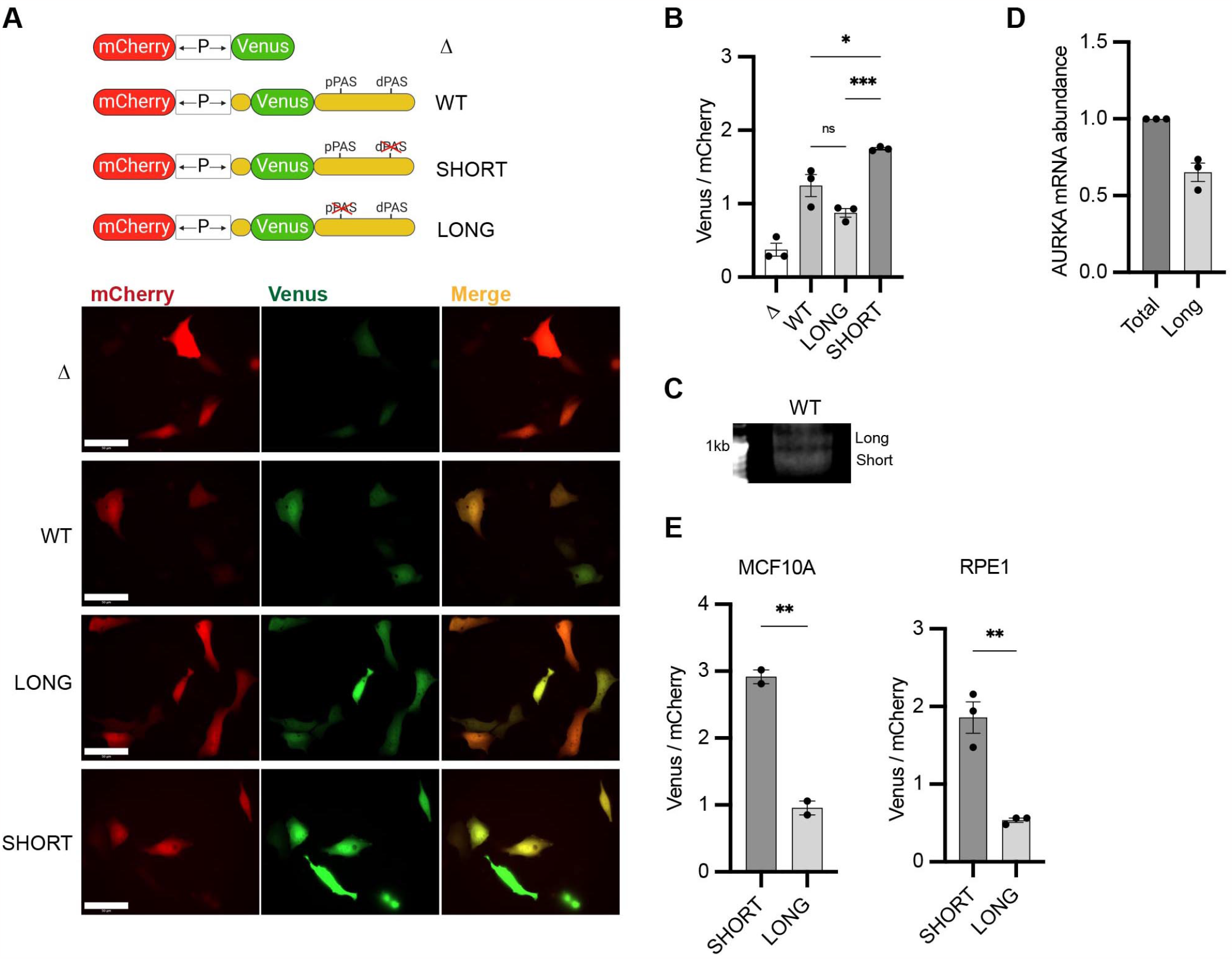
AURKA shows 3’UTR isoform-dependent protein expression. **(A)** *Top*. UTR-dependent protein expression reporters. Venus CDS is flanked by AURKA 5’UTR and 3’UTR, WT or PAS-mutated. *Bottom*. Representative snapshots of transfected U2OS. Scale bar 50μm. **(B)** Mean and s.e.m. of median Venus/mCherry MFI ratios from transfected U2OS from three biological replicates. n ≥ 129 cells per condition. Ordinary one-way ANOVA with Tukey’s multiple comparisons test. **(C)** 3’RACE of endogenous AURKA APA isoforms. **(D)** RT-qPCR of endogenous AURKA SLR in U2OS. Long isoform abundance plotted as fold change over total AURKA mRNA. 18S rRNA used as reference target. **(E)** Same as (B) but in MCF10A (*left*) and RPE1 (*right*) cells. n ≥ 55 cells per condition. Unpaired t-test. ns, not significant; *, p<0.05; **, p<0.01; ***, p<0.001.

We initially assessed the efficiency of the promoter bidirectionality. Correlation between Venus and mCherry expression was strongly maintained at the level of both fluorescence intensity (**Figure 2– figure supplement 1A**) and mRNA abundance (**Figure 1–figure supplement 1B**). Promoter strength was however not equal in both directions, since fewer copies of mCherry mRNA were transcribed compared to Venus mRNA (**Figure 2–figure supplement 1B**), despite mCherry fluorescence intensity being generally higher than that of Venus (**Figures 2A-B**, see Δ). We further assessed that mRNA of both mCherry and Venus was stable over time (**Figure 3B**, see Δ), and fluorescence of both proteins was stable over time and over different cell cycle stages (**Figure 2–figure supplement 1C-D**). Considering the short maturation time and long half-life of both Venus and mCherry proteins (55) (56), the assay allows to reliably measure effects of UTRs on reporter protein levels at any given time and regardless of cell cycle phases. As positive control of our assay, we recapitulated the higher protein expression from the short 3’UTR APA isoform of CDC6 mRNA, which has previously been observed (40) (**Figure 2–figure supplement 1E**).

**Figure 3:**
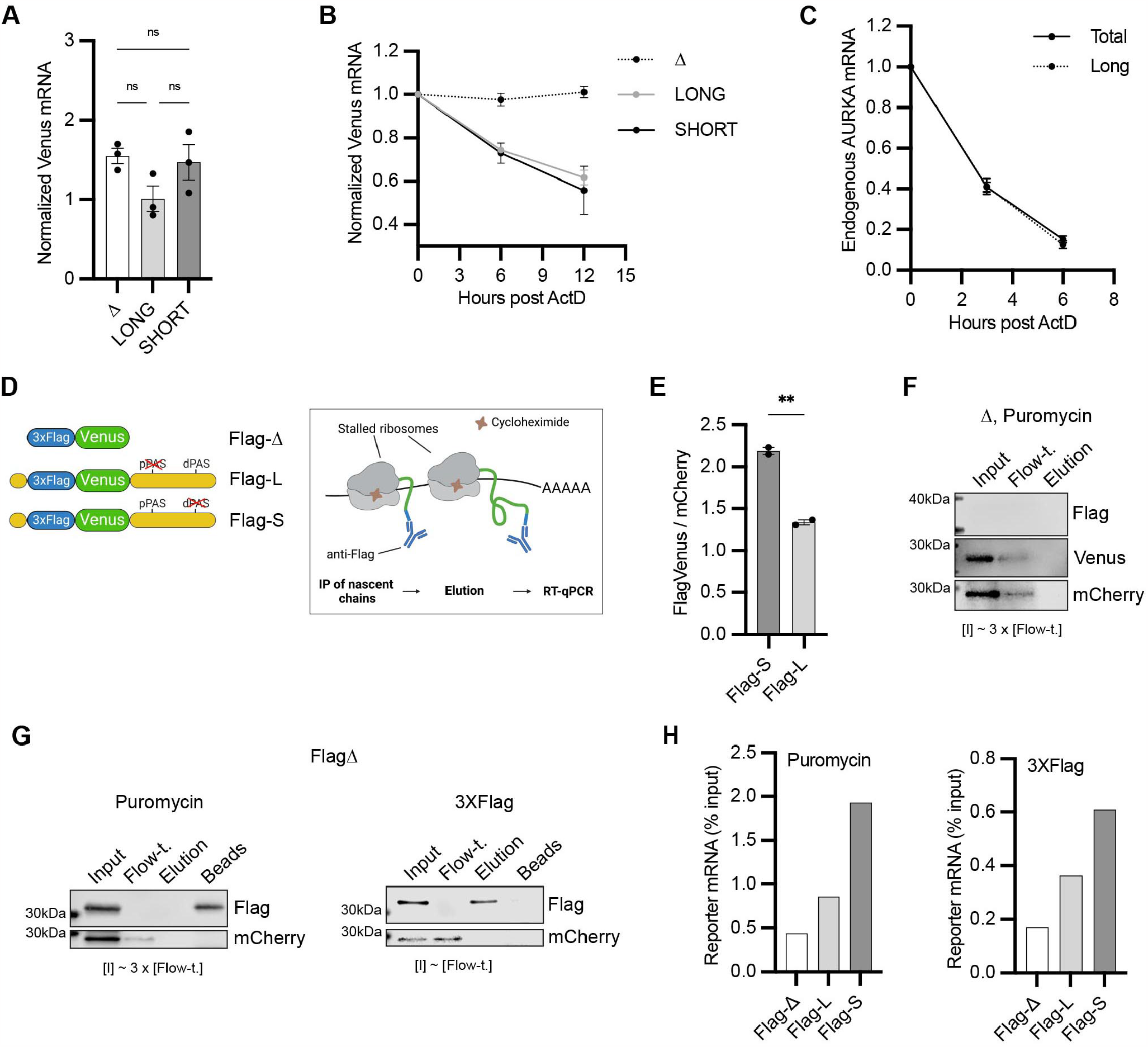
AURKA APA isoforms are translated with different efficiency. **(A)**,**(B)** RT-qPCR of reporter mRNAs abundance (A) and decay rate (B) from transfected U2OS. mCherry mRNA used as reference target. Ordinary one-way ANOVA with Tukey’s multiple comparisons test; ns, not significant. **(C)** Decay rate of endogenous AURKA mRNA as in (B). 18S rRNA used as reference target. Abundance of long isoform plotted as fold change over total AURKA mRNA. **(D)** Design of the Nascent Chain Immunoprecipitation (NC IP) reporters and assay. **(E)** Mean and s.e.m. of median FlagVenus/mCherry MFI ratios from transfected U2OS from two biological replicates. n ≥ 160 cells per condition. Unpaired t-test; **, p<0.005. **(F), (G)** Immunoblots of NC IP fractions using Δ (F) or Flag-Δ (G) reporter. mCherry used as negative control. **(H)** RT-qPCR of eluted reporter mRNAs. Results representative of three biological replicates.

Addition of AURKA UTRs to Venus CDS significantly increased protein expression (**Figures 2A-B**), likely due to the role of 5’UTR in facilitating translation (57). We found that the SHORT reporter generates significantly more protein compared to the LONG (**Figures 2A-B**). Moreover, similar protein expression levels from the WT and LONG reporters suggest that AURKA WT 3’UTR is processed with a preference for dPAS in U2OS cells. Accordingly, we could detect both endogenous AURKA APA isoforms in U2OS cells by 3’RACE (**Figure 2C**) and confirmed by RT-qPCR that AURKA long isoform is prevalent (∼60% of total AURKA mRNA) (**Figure 2D**). The different SLR observed between U2OS cells, normal breast tissues and TNBC cell lines and tissues (**Figures 1B-D**) indicates that AURKA 3’UTR isoform prevalence is dependent on cell type. In addition, the quantitative difference in reporter protein expression by the isoforms also varied among cell types, suggesting cell-specific regulation (**Figures 2B, E**, SHORT vs LONG). In sum, these results provide evidence for the first time of a role for APA in controlling AURKA protein expression.

### AURKA APA isoforms are translated with different efficiency

We next investigated the basis of the different protein expression between AURKA APA isoforms. Following transfection of U2OS cells with the constructs in **Figure 2A**, we first quantified the abundance of reporter mRNA isoforms (**Figure 3A**). We then assessed the isoforms decay rate by quantifying reporter mRNAs at multiple time points following arrest of transcription by Actinomycin D (ActD) (**Figure 3B**). We observed that while Venus mRNA lacking UTRs was highly stable, reporter mRNA levels decreased at faster rate when carrying UTRs (**Figure 3B**), indicating that the assay reports on UTR-dependent effects on mRNA stability. Both the abundance and stability of the SHORT and LONG reporter isoforms were similar (**Figures 3A-B**). We additionally found that the two endogenous AURKA 3’UTR isoforms also have similar decay rates, albeit decaying at a higher rate compared to the reporter mRNAs (**Figure 3C**), suggesting that features present in AURKA CDS might influence mRNA stability (58).

Because the reporter APA isoforms share similar abundance and stability, we wondered whether they undergo different translational regulation instead. To this aim, we adapted a biochemical translation efficiency (TE) assay from Williams *et al*. (59), which required addition of a 3XFlag tag to the N-terminus of Venus (FlagVenus) in our reporter constructs, and called this Nascent Chain Immunoprecipitation (NC IP) assay (**Figure 3D**). We first assessed that addition of the 3XFlag tag did not alter Venus expression (**Figure 3E**). In our NC IP assay, anti-Flag beads were used to immunoprecipitate nascent FlagVenus chains from ribosomes stalled by treatment with cycloheximide (CHX). Ribosome-mRNA complexes were eluted from the IP-immobilized nascent chains using puromycin, which causes release of nascent chains from ribosomes (60); RNA was then purified from the elution fraction and quantified by RT-qPCR to provide a measure of the amount of reporter mRNA undergoing translation. All fractions were then blotted for FlagVenus and mCherry (negative control) proteins to monitor their presence at different steps of the experiment. As expected, elution with puromycin retained FlagVenus on the beads, whereas mCherry, as well as untagged Venus, are lost in the flow-through (**Figure 3G**, *left*, **Figure 3F**). No reporter mRNA could be detected in the elution fraction when untagged Venus was used (data not shown). Alternatively, purified 3XFlag peptide was used to elute the nascent chain-ribosome-mRNA complexes from the beads following IP (**Figure 3G**, *right*). RT-qPCR quantification of the FlagVenus nascent chain-cognate reporter mRNAs revealed about twice more copies of Flag-S mRNA compared to Flag-L mRNA, regardless of the elution method (**Figure 3H**). This indicates that Flag-S mRNA is translated with higher efficiency than Flag-L mRNA. These results show that APA controls AURKA protein expression mainly via differential translational regulation of the 3’UTR isoforms.

### Translation rate of AURKA APA isoforms follows different cell cycle periodicity

Given the known cell cycle-dependent expression of AURKA (3), we tested whether differential translational efficiency of AURKA mRNA isoforms might contribute to this regulation. To avoid perturbation of translation provoked by classical cell synchronization methods (61), we used a live-cell fluorescence-based translation rate measurement assay in conjunction with a CDK2 activity sensor (62) for *in silico* cell cycle synchronization. We developed our assay of ‘Translation Rate Imaging by rate of Protein Stabilization’ (TRIPS) based on a previously introduced reporter system (63) (64). Our bidirectional promoter construct was modified to express superfolder GFP (sfGFP) fused to a mutated *E. coli* dihydrofolate reductase (DHFR-Y100I) destabilizer domain (DHFR-sfGFP), which is continuously degraded unless the stabilizer molecule trimethoprim (TMP) is added. Addition of TMP leads to an increase of sfGFP signal over time and, given the sfGFP short maturation time (65), the accumulation rate of sfGFP reflects DHFR-sfGFP protein synthesis rate (**Figure 4A**, *left*, **Figure 4–figure supplement 1A**). The ratio of the median of single-cell mCherry-normalized sfGFP signals at 2h to that at 0h of TMP treatment was therefore used as read-out for bulk translation rate. In accordance with our assay being designed to measure translation rate, sfGFP signal could not increase under TMP treatment in the presence of translation inhibitor CHX (**Figure 4B**). We also ensured that the increase in sfGFP signal was TMP-dependent (**Figure 4–figure supplement 1B**) and that TMP treatment affected neither mCherry expression (**Figure 4–figure supplement 1C**) nor DHFR-sfGFP mRNA abundance (**Figure 4C**). To probe translation rate at different cell cycle phases we used the CDK2 activity sensor (62) stably expressed in our U2OS cell line (U2OS^CDK2^) (**Figure 4–figure supplement 1A**) and called this assay ‘Cell cycle-dependent TRIPS’ (C-TRIPS) (**Figure 4A**, *right*).

**Figure 4:**
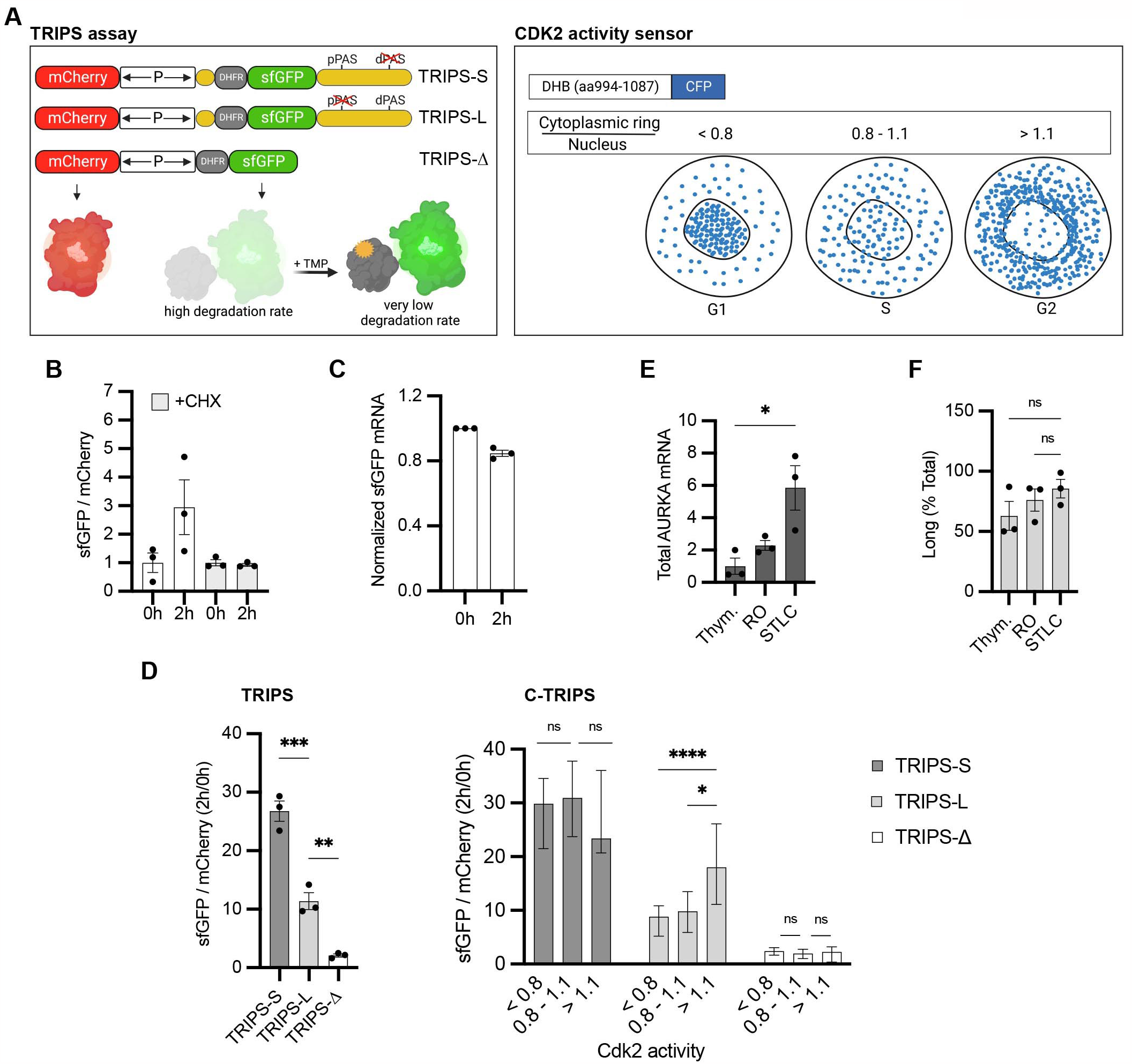
Translation rate of AURKA APA isoforms follows different cell cycle periodicity. **(A)** Design of the TRIPS reporters and assay (*left*) and CDK2 activity sensor (*right*). **(B)** Mean and s.e.m. of median sfGFP/mCherry MFI ratios from U2OS transfected with TRIPS-Δ and imaged at 0h and 2h of 50μM TMP treatment, with or without 0.1mg/ml CHX, from three biological replicates. Baseline at 0h. n ≥ 55 cells per condition. **(C)** RT-qPCR of sfGFP mRNA from U2OS transfected with TRIPS-Δ, at 0h and 2h of 50μM TMP treatment. mCherry mRNA used as reference target. RNA extracts at 0h of treatment used as reference sample. **(D)** TRIPS (*left*) and C-TRIPS (*right*) assays in transfected U2OS^CDK2^. n ≥ 200 cells per condition. *Left*. Mean and s.e.m. from three biological replicates. *Right*. Median and 95% CI of pooled data from *left*. Kruskal-Wallis with Dunnett’s multiple comparisons test. **(E)** RT-qPCR of endogenous AURKA mRNA in U2OS. 18S rRNA used as reference target. Baseline at G_1_/S. **(F)** Endogenous AURKA long isoform as in (E) plotted as percentage of total AURKA mRNA. **(D)** *Left*, **(E), (F)** Ordinary one-way ANOVA with Tukey’s multiple comparisons test. ns, not significant; *, p<0.05; **, p<0.01; ***, p<0.0005; ****, p<0.0001.

To test the translation rate of the individual AURKA APA isoforms, we flanked DHFR-sfGFP CDS with AURKA PAS-mutated UTRs (TRIPS-L, TRIPS-S) (**Figure 4A**, *left*). We found that our TRIPS assay could recapitulate the difference in translation efficiency of the isoforms previously observed (**Figures 3H, 4D**, *left*). Importantly, expression of the CDK2 activity sensor did not affect cellular translation, as the different translation rate of AURKA 3’UTR isoforms could be reproduced in U2OS cells lacking the sensor (**Figure 4–figure supplement 1D**). Following measurements of bulk translation rates (**Figure 4D**, *left*), we then binned single-cell translation rate values into three intervals of CDK2 activity (**Figure 4D**, *right*). Results of our C-TRIPS assay revealed that, while TRIPS-Δ is translated constantly during the cell cycle, translation rate of TRIPS-L is regulated in the cell cycle. This isoform showed lower translation rate in G_1_ and S and an enhanced rate at G_2_, consistent with the increase in both AURKA mRNA and protein levels that occurs in preparation for mitosis (3). By contrast, TRIPS-S was translated constantly through the cell cycle and at a maximal rate already in G_1_ (**Figure 4D**, *right*), indicating that this isoform is insensitive to cell cycle regulation of AURKA translation rate.

Furthermore, we quantified abundance of endogenous AURKA APA isoforms at different stages of the cell cycle, by performing RT-qPCR following synchronization in G_1_/S, G_2_, or M phases (**Figure 4E**). The expected changes in AURKA mRNA abundance following each treatment represent a positive control for the synchronization. However, abundance of the long isoform changed quite concomitantly with changes in total AURKA mRNA levels (**Figure 4F**). This suggests that the same ratio of 3’UTR isoforms is rather maintained throughout the cell cycle and that AURKA APA is not cell cycle regulated. These results not only provide strong, independent validation of our finding that elements present in 5’ and 3’ UTR of AURKA enable translational activation but additionally indicate that elements present on the long 3’UTR might account for its different pattern of translation during interphase, as lack of these on the short 3’UTR allow escape from cell cycle phase-dependent translation.

### Translational periodicity of long 3’UTR isoform is regulated by *hsa-let-7a* miRNA

Among known post-transcriptional regulators, miRNAs are widely recognized as molecular regulators of both mRNA stability and translation (66). We interrogated miRDB (mirdb.org) (67) to search for miRNAs that could be involved in the differential regulation of the two AURKA mRNA isoforms, and selected *hsa-let-7a* miRNA given its widely established tumor-suppressor role of in TNBC. We assessed the *hsa-let-7a* targeting of AURKA 3’UTR by co-transfecting our AURKA UTR-dependent protein expression reporters (**Figures 2A**) with *hsa-let-7a* or a negative control miRNA that does not have any target in the human genome. As positive control of the assay, we cloned Myeloid Zinc Finger1 (MZF1) 3’UTR downstream Venus CDS in our Δ reporter and could reproduce the previously reported targeting of MZF1 3’UTR by *hsa-let-7a* (68) (**Figure 5B**). Protein expression from the LONG reporter mRNA was reduced by *hsa-let-7a*, whereas that from the SHORT mRNA and from a LONG mRNA that lacks the *hsa-let-7a* binding site (Δlet7a) was not (**Figure 5B**). Also, the loss of *hsa-let-7a* targeting was sufficient to increase protein expression from the LONG reporter mRNA (compare LONG + NC vs Δlet7a + NC). To confirm that altered expression was due to the lack of *hsa-let-7a* targeting and not an effect of the mutation itself, we also observed an increase in protein expression when we co-transfected our LONG reporter and an inhibitor of *hsa-let-7a* (*anti-let7a*) (**Figure 5C**).

**Figure 5:**
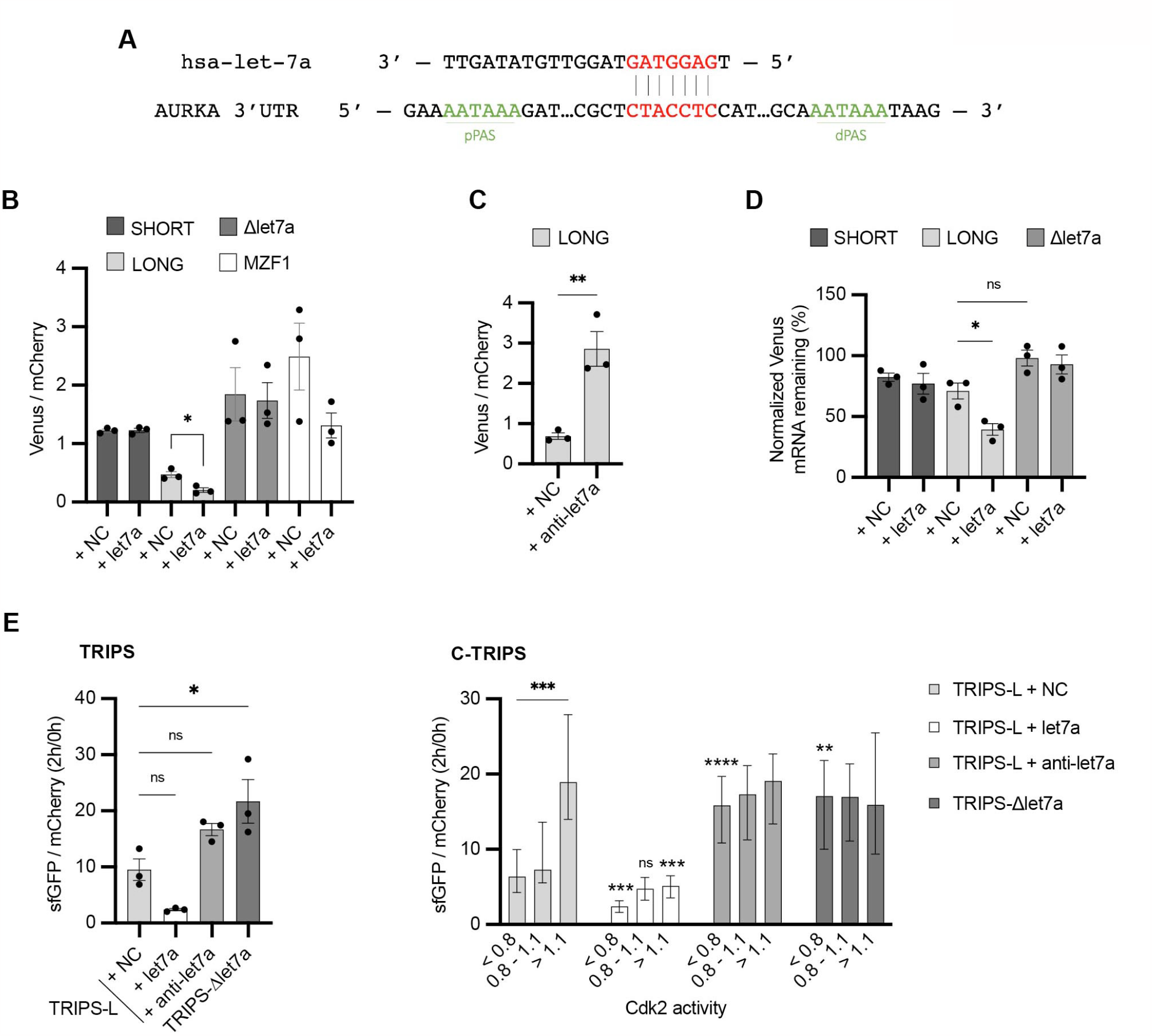
Translational periodicity of long 3’UTR isoform is regulated by *hsa-let-7a* miRNA. **(A)** Complementarity of *hsa-let-7a* binding to AURKA 3’UTR. **(B)** Mean and s.e.m. of median Venus/mCherry MFI ratios from U2OS co-transfected with 250nM *hsa-let-7a* or a negative control (NC) miRNA from three biological replicates. n ≥ 182 cells per condition. Unpaired t-test. **(C)** Same as (B) but co-transfecting 300nM *anti-let-7a* or NC. n ≥ 94 cells per condition. Unpaired t-test. **(D)** RT-qPCR of reporter mRNAs abundance from U2OS cells transfected as (B), at 8h of 10μg/ml ActD. mCherry mRNA used as reference target. Ordinary one-way ANOVA with Tukey’s multiple comparisons test. **(E)** TRIPS (*left*) and C-TRIPS (*right*) assays in transfected U2OS^CDK2^. n ≥ 162 cells per condition *Left*. Mean and s.e.m. from three biological replicates. Ordinary one-way ANOVA with Dunnett’s multiple comparisons test vs NC. *Right*. Median and 95% CI of pooled data from *left*. Kruskal-Wallis with Dunnett’s multiple comparisons test vs NC of the respective phase. ns, not significant; *, p<0.05; **, p<0.01; ***, p<0.001; ****, p<0.0001.

In order to assess the role of *hsa-let-7a* in controlling decay rate of the target mRNA, we next co-transfected our Venus reporters and either *hsa-let-7a* or negative control miRNA and quantified reporter mRNA abundance after 8h of ActD treatment. We found that stability of the LONG reporter mRNA was significantly reduced by *hsa-let-7a*, whereas that of the SHORT reporter mRNA was unaltered. Additionally, mutation of the *hsa-let-7a* binding site slightly increased reporter mRNA stability (compare LONG + NC vs Δlet7a + NC) (**Figure 5D**).

We then performed our C-TRIPS assay co-transfecting the TRIPS-L reporter and *hsa-let-7a* or control miRNA and found that *hsa-let-7a* reduced both bulk translation rate (**Figure 5E**, *left*) and translation rate at all interphase stages (**Figure 5E**, *right*) of the long 3’UTR. Furthermore, we asked whether loss of *hsa-let-7a* targeting is sufficient to cause loss of translational regulation of the long isoform during the cell cycle. For this, we performed the C-TRIPS assay using a TRIPS-L reporter with mutations in the *hsa-let-7a* binding site (TRIPS-Δlet7a) or, alternatively, co-transfecting the TRIPS-L reporter and the *hsa-let-7a* inhibitor *anti-let7a*. Interestingly, in both cases, loss of *hsa-let-7a* targeting only increased translation rate in G_1_ and S, but not G_2_ (**Figure 5E**, *right*), suggesting that the targeting in G_2_ is not likely to occur unless in conditions of excess *hsa-let-7a*.

In conclusion, our results show that *hsa-let-7a* only silences AURKA long 3’UTR isoform by both promoting mRNA degradation and reducing translation rate, and that *hsa-let-7a* targeting is responsible for the cell cycle-dependent translational regulation of AURKA long 3’UTR isoform.

### Increased AURKA short/long ratio is sufficient to disrupt cell behavior

Having established that APA plays a role in regulating AURKA expression, we tested the idea that AURKA APA directly contributes to cancer cell behaviour, by performing genome editing to alter AURKA APA in wild-type U2OS cells. We used Cas9^D10A^-mediated double-nicking strategy and mutated the endogenous dPAS on *AURKA* 3’UTR with the aim of silencing expression of AURKA long 3’UTR isoform (**Figure 6A**). Two mutant clones with disrupted dPAS site were obtained (ΔdPAS#1, ΔdPAS#2) (**Figure 6–figure supplement 1A**) and were used for subsequent functional analyses. Qualitative assessment of AURKA 3’UTR isoforms ratio in these clones by 3’RACE found the long 3’UTR isoform to be undetectable in the mutated cell lines (**Figure 6B**), indicating that the genetic editing successfully prevents usage of the dPAS site for cleavage and polyadenylation.

**Figure 6:**
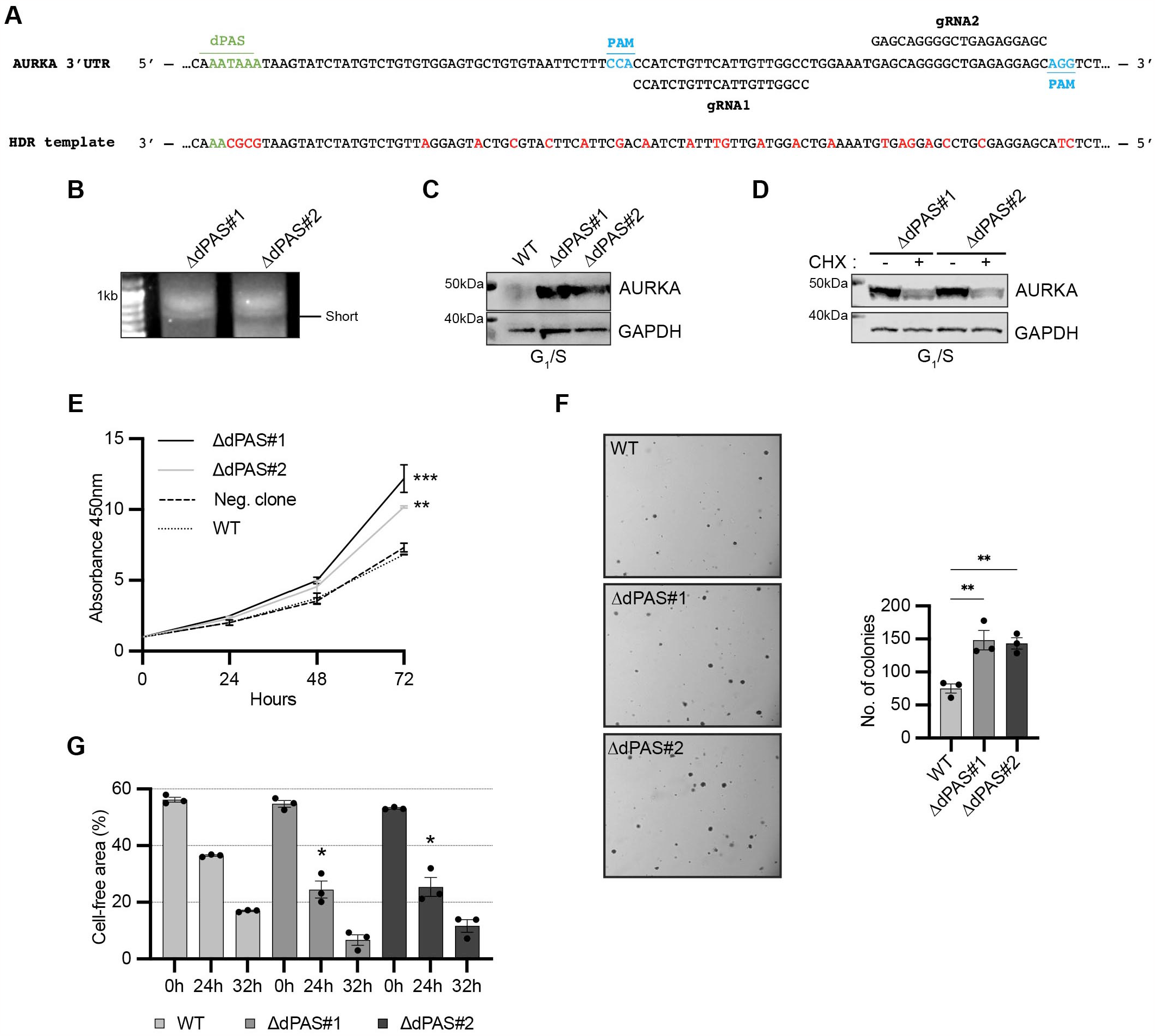
Increased AURKA short/long ratio is sufficient to disrupt cell behavior. **(A)** Design of CRISPR editing. Nucleotide substitutions in red. **(B)** 3’RACE of endogenous AURKA APA isoforms. **(C)**,**(D)** Western blot after G_1_/S enrichment (C) and with or without 6h treatment with 0.1mg/ml CHX (D). Blots representative of three biological replicates. **(E)** CCK8 assay. A non-parental WT U2OS cell line used as negative control. Ordinary one-way ANOVA with Dunnett’s multiple comparisons test vs WT; **, p<0.005; ***, p<0.001. **(F)** *Left*. Representative images of cells grown in soft agar. *Right*. Mean number of clones and s.e.m. of three biological replicates. **(G)** Measurement of migration rate. **(F), (G)** Ordinary one-way ANOVA with Dunnett’s multiple comparisons test vs WT; *, p<0.05; **, p<0.01.

We then examined expression of AURKA in the mutated cell lines by immunoblot of extracts from cell populations enriched for the G_1_/S phase of the cell cycle, where AURKA expression is the lowest in unmodified cells. We observed AURKA was expressed at higher levels in ΔdPAS#1 and ΔdPAS#2 cells compared to WT cells (**Figure 6C, Figure 6–figure supplement 1B**). AURKA expression in G_1_/S was reduced in the mutated cell lines when treated with CHX, indicating that translation of the short isoform is active in this phase (**Figure 6D**). Because AURKA overexpression is a common feature of cancer, we interrogated the mutated cell lines for changes in cancer-relevant behaviour. Consistent with a role of AURKA overexpression in accelerating the cell cycle and favoring cell proliferation, we found a higher rate of proliferation of ΔdPAS#1 and ΔdPAS#2 cells compared to WT cells using the CCK8 assay to measure metabolic activity (**Figure 6E**). Additionally, we assessed the ability of anchorage-independent growth, which closely correlates with tumorigenicity in animal models, by growing cells in soft agar. ΔdPAS#1 and ΔdPAS#2 cells resulted more capable to survive and grow in the absence of anchorage to their neighboring cells compared to WT cells (**Figure 6F**). AURKA also regulates organization of microtubules required for cellular migration and it also enhances migration of tumor cells through several pathways. For example, AURKA activates the Cofilin-F-Actin pathway leading to breast cancer metastases (1). We were therefore prompted to assess the motility of ΔdPAS#1 and ΔdPAS#2 cells in a 2D cell migration assay. Our result shows a higher rate of migration of ΔdPAS#1 and ΔdPAS#2 cells compared to WT cells (**Figure 6G**).

In summary, our results show that AURKA overexpression caused by a disruption in the SLR ratio of APA isoforms in favor of the short isoform contributes to cancer-like cell behavior.

## DISCUSSION

In this study, we describe for the first time a molecular mechanism involving the dysregulation of APA and the differential targeting of AURKA APA isoforms by *hsa-let7-7a*, a tumor suppressor miRNA, that is sufficient for the oncogenic activation of AURKA at the post-transcriptional level. We also shed light on the cell cycle-dependent regulation of AURKA translation and introduce novel and improved methods for measurements of post-transcriptional gene expression of individual mRNAs of interest.

As a consequence of tandem 3’UTR APA, a short and a long 3’UTR isoform are generated for AURKA mRNA. The short/long ratio of these isoforms is cell-type defined and is not dependent on the cell cycle, indicating that cell-type specific/cell cycle-independent factors are involved in establishing the short/long ratio. This is unexpected given that periodic regulation of APA may be a characteristic of many cell cycle genes (69). Because protein expression differs from the two isoforms, AURKA short/long ratio is a crucial element defining AURKA expression levels. Ours and other studies detected increased short/long ratio of AURKA APA isoforms in TNBC and found this correlates with worse disease-free survival (14) (35). Here we reveal insights into the molecular basis for the correlation, by showing that the increased and cell cycle-independent translation rate of the short isoform can lead to marked overexpression of AURKA in interphase. Our results support the hypothesis that deregulation of expression by disruption of APA is sufficient to drive AURKA oncogenic properties, such as promoting increased proliferation and migration rate. Whether this is also sufficient to drive cancer-cell transformation remains to be explored. It has been known for many years that overexpression of AURKA induces mitotic defects, aneuploidy, as well as acceleration of the cell cycle, epithelial-to-mesenchymal transition and migration (1) (70), whilst the cellular background for AURKA’s transforming potential is also of importance (71). Nonetheless, the significance of AURKA overexpression specifically in G_1_ for the exertion of potential oncogenic functions in this phase represents an emerging field of research (2) (44) (72). It is possible that normal AURKA functions are exerted at low levels of expression in G_1_ become oncogenic at high levels of expression. This would not be surprising giving how the roles played by AURKA in G_1_ revolve around regulation of transcription, mitochondria fitness and cellular metabolism.

Our work does not research the cause of the disrupted APA in TNBC. This could be concomitant to a global 3’UTR shortening, for example due to altered expression in cancer of C/P factors (9) (14), or of their regulators (21), or even altered RNA Pol II elongation dynamics (73). However, these phenomena have not been extensively explored in TNBC (74). Alternatively, disrupted APA could represent an AURKA gene-specific phenomenon due to presence of Single Nucleotide Polymorphisms (SNPs) on the 3’UTR, either on the C/P site or in proximity, that could affect PAS choice by the C/P machinery. Finally, a mRNA-dependent mechanism whereby the short 3’UTR isoform itself regulates AURKA protein function or localization throughout or at specific phases of the cell cycle, licensing oncogenic advantage, should not be excluded (73).

The periodicity of AURKA expression is an important requirement for correct progression through the cell cycle. For example, ubiquitin-mediated proteolysis is a critical pathway for irreversible AURKA inactivation, which must occur for ordered transition to interphase following mitosis (44) (75). One important but unanswered question is whether and to what extent translation regulation accounts for the increase and decrease of AURKA expression during the cell cycle. Implementation of our TRIPS assay, which measures protein synthesis rate independent of changes in mRNA abundance, allowed detection of active regulatory mechanisms of translation occurring at different cell cycle phases. This new evidence integrates well with the notion that transcriptional, post-transcriptional, and post-translational mechanisms all combine to provide *AURKA* gene with a characteristic pattern of expression (3). Our work shows that translation of AURKA is regulated by *hsa-let-7a* miRNA. Because we could still detect active protein synthesis from the long isoform under *hsa-let-7a* overexpression using our TRIPS assay, and we could also immunoprecipitate tagged nascent chains from the long isoform in U2OS cells, where *hsa-let-7a* is expressed, it is unlikely that *hsa-let-7a* blocks translation at the level of initiation, but it rather slows down the rate of translation elongation. This is in accordance with a study showing that *hsa-let-7a* co-sediments with actively translating polyribosomes (76), a generally proved mechanism of miRNA action (77). In addition, because we observed that *hsa-let-7a* can also control the decay rate of the long isoform, it is possible that a reduction in translation elongation rate may be required to mediate degradation of the mRNA (78). We also show that the differential *hsa-let-7a* targeting through the cell cycle is a mechanism responsible for AURKA periodic translational control. Based on evidence from our C-TRIPS assays to assess the temporal *hsa-let-7a* targeting of the long isoform at different phases of the cell cycle, and on the published evidence that *hsa-let-7a* levels are constant during the cell cycle of human cancer cells as well as untransformed fibroblasts (79), we propose that: (i) *hsa-let-7a* targeting is productive in G_1_ and S, and is therefore responsible for the low AURKA translation rate in these phases; (ii) the targeting is not occurring in G_2_ phase, except in excess of *hsa-let-7a*, possibly because *hsa-let-7a* overexpression saturates sequestering factors that prevent its binding to the long isoform. Further investigations will be required to understand this mechanism in more detail.

The characterization of gene-specific post-transcriptional dynamics is desirable for a complete understanding of gene expression regulation. Here, we have developed transient single-cell and biochemical assays to rapidly study mRNA-specific gene expression in a way that measures post-transcriptional events exclusively. Importantly, the assays can be used to test the effect of regulators such as therapeutic miRNAs or drugs on protein expression, mRNA processing, and translation of selected genes.

In conclusion, our study reveals a strong cooperation between APA and miRNA targeting in controlling gene expression dynamics of AURKA and its oncogenic potential. It also provides a workflow to assess the role of mRNA-specific post-transcriptional processing and regulators. Our work additionally highlights a molecular mechanism that could represent an actionable target of RNA-based therapeutics.

## MATERIALS AND METHODS

### *In silico* analysis of APA in TNBC

Publicly available CEL files and associated metadata of microarray results were downloaded from NCBI Gene Expression Omnibus (GEO) repository. APADetect tool (35) was used to detect and quantify APA events in TNBC patients and normal breast tissue. CEL files of Human Genome U133A (HGU133A, GPL96) and U133 Plus 2.0 Arrays (HGU133Plus2, GPL570) were analyzed to identify intensities of probes that were grouped based on poly(A) site locations extracted from PolyA_DB (38). Mean signal intensities of proximal and distal probe sets for AURKA were calculated and used as indicators of ‘Short’ and ‘Long’ AURKA 3’-UTR isoforms’ abundance. The ratio of proximal probe set mean (S) to the distal probe set (L) defined as short to long ratio (SLR). SLR values were subjected to significance analysis of microarrays (SAM), as implemented by the TM4 Multiple Array Viewer tool (39), for statistical significance after log normalization.

### Molecular cloning

The following UTR sequences were obtained by gene synthesis (Genewiz from Azenta Life Sciences, European Genomics Headquarters, Germany): AURKA WT 5’UTR and 3’UTR (769bp) (NM_003600.4), MZF1 3’UTR (NM_003422.3), AURKA individual PAS-mutated 3’UTRs (AATAAA>AATCCC). CDC6 3’UTRs were from (40). For *Δ* reporter, mCherry and Venus ORFs were inserted into the MCSs of *pBI-CMV1* (631630, Clontech, TakaraBio). *WT, SHORT* and *LONG* reporters were generated by assembly (NEBuilder® HiFi DNA Assembly Cloning Kit, E5520S, NEB) of AURKA 5’UTR, Venus ORF and AURKA wt, short (dPAS mutated) or long (pPAS mutated) 3’UTR, and insertion into *pBI-CMV1-mCherry. CDC6_L, CDC6_S*, and *MZF1* reporters were generated by insertion of CDC6 long and short 3’UTRs, and MZF1 3’UTR downstream Venus CDS in Δ reporter. *LONG-Δlet7a* was generated by site-directed mutagenesis of *LONG* reporter with the following forward and reverse primers: 5’-CACGCACCATTTAGGGATTTGCTTG-3’ and 5’AGCACGTGTTCCTA-TTTTTCACACTC-3’. *Flag-S* and *Flag-L* reporters were generated by assembly of AURKA 5’UTR, 3XFlag-Venus ORF and AURKA short (dPAS mutated) or long (pPAS mutated) 3’UTR, and insertion into *pBI-CMV1-mCherry. Flag-Δ* was generated by insertion of 3XFlag-Venus ORF into *pBI-CMV1-mCherry*. For *TRIPS* reporters, the DHFR-sfGFP ORF was PCR amplified from *pHR-DHFRY100I-sfGFP-NLS-P2A-NLS-mCherry-P2A_Emi1 5’ and 3’UTR* plasmid, a gift from Ron Vale (Addgene plasmid #67930), and inserted into *pBI-CMV1-mCherry* (*TRIPS-Δ)* or assembled with AURKA 5’UTR and AURKA wt 3’UTR (*TRIPS-WT*), short 3’UTR (*TRIPS-S*) or long 3’UTR (*TRIPS-L*) before insertion. *TRIPS-Δlet7a* was generated by site-directed mutagenesis of *TRIPS-L* reporter with the primers above. NEB® 5-alpha Competent *E. coli* (High Efficiency) (C2987I, NEB) was used.

### Cell lines and drug treatments

Human U2OS, U2OS^CDK2^, BT20, HCC1143, HCC1937, MDA-MB-157, MDA-MB-231 and MDA-MB-468 cell lines were cultured in DMEM (ThermoFisher) supplemented with 10% FBS (Sigma), 200μM GlutaMAX-1 (ThermoFisher), 100U/ml penicillin, 100μg/ml streptomycin, and 250ng/ml fungizone at 37°C with 5% CO_2_. U2OS^CDK2^ cells cultures were supplemented with 500μg/ml G-418 (Sigma). Human MCF10A were cultured in filtered DMEM-F12 (ThermoFisher) supplemented with 10% FBS, 20ng/ml EGF (Peprotech), 0.5mg/ml hydrocortisone (Sigma), 100ng/ml cholera toxin (Sigma), 10μg/ml insulin (Sigma), 100U/ml penicillin, 100μg/ml streptomycin, and 250ng/ml fungizone at 37°C with 5% CO_2_. Human RPE1 cells were cultured as previously described (41). Breast cancer cell lines were purchased from DSMZ (Germany) or ATCC (USA) with authentification certificates including STR profiling; all cell lines used in the study were free of mycoplasma contamination. Cell populations were enriched for G_1_/S phase by incubating with 2.5mM thymidine for 24h, for G_2_ phase by incubating with 10μM RO3306 (4181, Tocris Bioscience) for 16h, for M phase by incubating with 10μM S-trityl l-cysteine (STLC) (2191/50, Tocris Bioscience) for 16h and mitotic cells were then collected by shake-off. CHX (239763, Sigma), TMP (92131, Sigma), and DMSO (sc-358801, Insight biotech) were used as indicated in figure legends.

### Transfections

U2OS and RPE1 cells (5×10^6^) were electroporated (Neon Transfection System, Invitrogen) using 1150V pulse voltage, 30ms pulse width, and 2 pulses. U2OS^CDK2^ and MCF10A cells (4×10^4^) were transfected using Lipofectamine 3000 Transfection Reagent (L3000001, ThermoFisher) according to the manufacturer’s instructions. MISSION® microRNA Mimic *hsa-let-7a* (HMI0003, Sigma), miRNA Mimic Negative Control (ABM-MCH00000, abm), and Anti-miR™ miRNA Inhibitor (AM17000, ThermoFisher) were co-transfected by Lipofectamine RNAiMAX Transfection Reagent (13778100, ThermoFisher) according to the manufacturer’s instructions. All analyses were carried out 24h post transfection.

### Live-cell fluorescence microscopy

Live-cell microscopy was performed using Olympus IX81 motorized inverted microscope, Orca CCD camera (Hamamatsu Photonics, Japan), motorized stage (Prior Scientific, Cambridge, UK) and 37°C incubation chamber (Solent Scientific, Segensworth, UK) fitted with appropriate filter sets and a 40X NA 1.42 oil objective. Images were collected in the 490nm (Venus, sfGFP), 550nm (mCherry) and 435nm (CFP) channels using Micro-Manager software (42). Image analysis was performed using a customized plug-in tool (Andrew Ying, 2017) in Image J (43), which calculates mean fluorescence intensity (MFI) by measuring average, background-subtracted grey values over regions of interest (ROIs) of defined diameter around manually selected points in the cell.

### Western blot

Western blot was performed as previously described (44). PageRuler Prestained Protein Ladder (26616, ThermoFisher) was used. Primary antibodies used: mouse anti-AURKA mAb (1:1000; Clone 4/IAK1, BD Transduction Laboratories), rabbit anti-GFP (1:5000; Ab290, Abcam), rabbit anti-GAPDH (1:4000; 2118S, CST), mouse anti-mCherry (1:1000; ab167453, Abcam), mouse ANTI-FLAG® M2 mAb (1:1000; F1804, Sigma). Secondary antibodies were rabbit (P044801-2) or mouse (P044701-2) HRP-conjugated (Dako, Agilent), used at 1:10000 dilution, and detection was performed via Immobilon Western Chemiluminescent HRP Substrate (WBKLS0100, Millipore) on an Odyssey Fc Dual-Mode Imaging System (LICOR Biosciences). Raw images of blots are shown in **Supplementary File 1**.

### RNA extraction and RT-qPCR

RNA extracts were collected using Total RNA miniprep kit (T2010S, NEB). DNA was in-column digested with DNase I. Aliquots were stored with 5μM EDTA at -20°C for a week or flash-frozen in dry ice and transferred at -80°C. A micro-volume spectrophotometer (NanoDrop™ Lite, VWR) was used to assess A_260_/A_280_ ratios of ∼ 2.0 and A_260_/A_230_ ratios of 2.0-2.2. DNA contamination was also tested (**Supplementary File 2**). RT-qPCR was performed using Luna® Universal One-Step RT-qPCR Kit (E3005S, NEB). 20μl reactions using 200nM primers and <100ng RNA were run on ABI StepOnePlus Real Time PCR system following manufacturer’s instructions. Primers were designed at Eurofins Genomics. ΔC_t_ or ΔΔC_t_ method was used for relative quantifications accounting for the primer pairs amplification efficiency. Three technical replicates were performed in each biological replicate. Results shown as mean and s.e.m. of three biological replicates. Sequences of primers, validation of amplification efficiency of primer pairs, and checklist of MIQE guidelines (45) are shown in **Supplementary File 2**. Microsoft Excel and GraphPad Prism were used for data analysis.

### mRNA decay measurement

Cells were treated with 10μg/ml Actinomycin D (ActD) (10043673, Fisher Scientific) and RNA was isolated from cells at the indicated time points after inhibition of transcription. Target mRNA was quantified by RT-qPCR using the ΔΔC_t_ method with indicated reference targets and corresponding RNA extracts at 0h of ActD treatment as reference sample. Mean and s.e.m. of three biological replicates are shown at each time point.

### Nascent Chain Immunoprecipitation

Transfected cells were treated with 0.1mg/ml CHX for 15 minutes, then washed, centrifuged, and resuspended in ice-cold lysis buffer [100mM Tris-HCl pH 7.5, 500mM LiCl, 10mM EDTA, 0.1mg/ml CHX, 0.1% Triton X-100, 100U/ml RNasin (3335399001, Merck) and cOmplete EDTA-free protease inhibitor cocktail (11836170001, Roche)] and incubated 15 minutes on ice. Lysates were cleared by centrifugation and supernatant was incubated with anti-Flag M2 magnetic beads (M8823, Sigma) overnight at 4 °C rotating. The bound fraction was washed twice (10mM Tris-HCl pH 7.5, 600mM LiCl, 1mM EDTA, 100U/ml RNasin, 0.1mg/ml CHX). Followed elution with 10mM Tris-HCl pH 7.5, 600mM LiCl, 1mM EDTA, 100U/ml RNAsin, 0.1mg/ml Puromycin (J67236.XF, Alfa Aesar), or with 3XFLAG peptide buffer (F4799, Sigma), for 30 minutes rotating at 4°C. RNA was purified (Monarch® RNA Cleanup Kit, T2040L, NEB) from fractions and samples were stored as above. Aliquots of each fraction were mixed 1:1 with NuPAGE® LDS Sample Buffer 4X (NP0007, Invitrogen), boiled 3 minutes at 95°C and stored at -20°C.

### 3’RACE

cDNA synthesis was performed using the Transcriptor Reverse Transcriptase (3531317001, Roche) with an oligo-dT anchor primer (5’-GACCACGCGTATCGATGTCGAC-TTTTTTTTTTTTTTTTV-3’). In the first round of PCR, an AURKA-specific forward primer (5’-TCCATCTTCCAGGAGGACCACTCTCTG-3’) was used with a reverse primer for the oligo-dT anchor sequence (Anchor_R: 5’-GACCACGCGTATCGATGTCGAC-3’). In the second round of PCR (nested), a new AURKA-specific forward primer (5’-CGGGATCCATATCACGGGTTGAATTCACATTC-3’) was used with Anchor_R. Nested PCR was performed using a 1:10 dilution of first PCR product as template. PCR product was visualized in agarose gels and imaged as above.

### Generation of ΔdPAS cell lines

Two guide RNAs (gRNA1: 5′-GGCCAACAATGAACAGATGG-3′ and gRNA2: 5′-GAGCAGGGGCTGAGAGGAGC-3′) were cloned into AIO-GFP (46), a gift from Steve Jackson (Addgene plasmid #74119). The donor DNA template for homology-directed repair (HDR) was cloned into a separate vector expressing mRuby (mRuby-HDR). AIO-GFP-gRNAs and mRuby-HDR were co-transfected into U2OS cells and 48h after transfection GFP^+^mRuby^+^ cells were sorted at single-cell density into multiple 96-well plates for clonal expansion. Cell populations were individually screened for dPAS mutation by touch-down PCR. Mutants were confirmed by Sanger sequencing of the genomic locus.

### Cell Counting Kit-8 (CCK-8) assay

Cells were seeded into 96-well plates at a density of 4000 cells/well. CCK-8 (96992, Merck) was used according to the manufacturer’s instructions and measurements were performed at the indicated time points. The OD_450_ value was determined using a CLARIOstar Plus microplate reader (BMG LABTECH). Mean and s.e.m. of three biological replicates shown for each time point.

### Colony formation assay in soft agar

A 0.6% base agarose was prepared in 6-well plates and cells were seeded at density of 15,000 cells per well prior mixing with agarose to a final agarose concentration of 0.3%. Fresh medium was added every 3 days. Colonies were imaged and counted after 10 days using phase contrast microscopy under 4X magnification.

### Migration assay

A suspension of 50,000 cells was added to each well of a Culture-Insert 2 Well (IB-80209,Thistle Scientific Ltd) and grown into a monolayer. After insert removal, cells were washed and serum-free medium was added. At indicated time points, images of three different fields were acquired for every condition, using phase contrast microscopy under 10X magnification. The % of cell-free area was calculated using Wound_healing_size_tool plugin (47) in ImageJ and is shown as mean of three fields and s.e.m. for three biological replicates.

### Statistical analyses

Graph Pad Prism 9 software (version 9.5.0) (Graph Pad Software Inc.) was used to generate graphs and perform statistical analyses. Statistical parameters including the sample size, the statistical test used, statistical significance (p value), and the number of biological replicates are reported in figure legends or methods.

### Materials availability

Materials from this study are available from the corresponding author CL upon reasonable request.

## DATA AVAILABILITY

The data underlying this article are available in the article and in its online supplementary material.

## FUNDING

This work was supported by Biotechnology and Biological Sciences Research Council (BBSRC) [grant no. BB/R004137/1] to CL. RC is supported by David James Studentship from the Department of Pharmacology.

## ACKNOWLEDGEMENTS

We thank past and present members of Lindon Lab for enriching discussions throughout the study. We are grateful to Chiara Marcozzi for advice on CRISPR/Cas9 and James Adeosun for help with molecular cloning. We are also grateful to Tim Weil, Adrien Rousseau, and Francesco Nicassio for insightful comments. Cartoon figures were created using BioRender.com.

## CONFLICT OF INTEREST

None declared.

## AUTHOR CONTRIBUTIONS

Conceptualization – RC, HBA, EEB, CL

Methodology – RC, HBA

Software – TT, HBA

Validation – HBA, RC

Formal analysis – RC, HBA

Investigation – RC, HBA

Resources – RC, HBA, EEB, CL

Data curation – RC

Writing – original draft preparation – RC

Writing – review & editing – RC, HBA, CL

Visualization – RC

Supervision – CL, EEB

Project administration – RC, CL

Funding acquisition – CL

---

**Figure 2—figure supplement 1.**
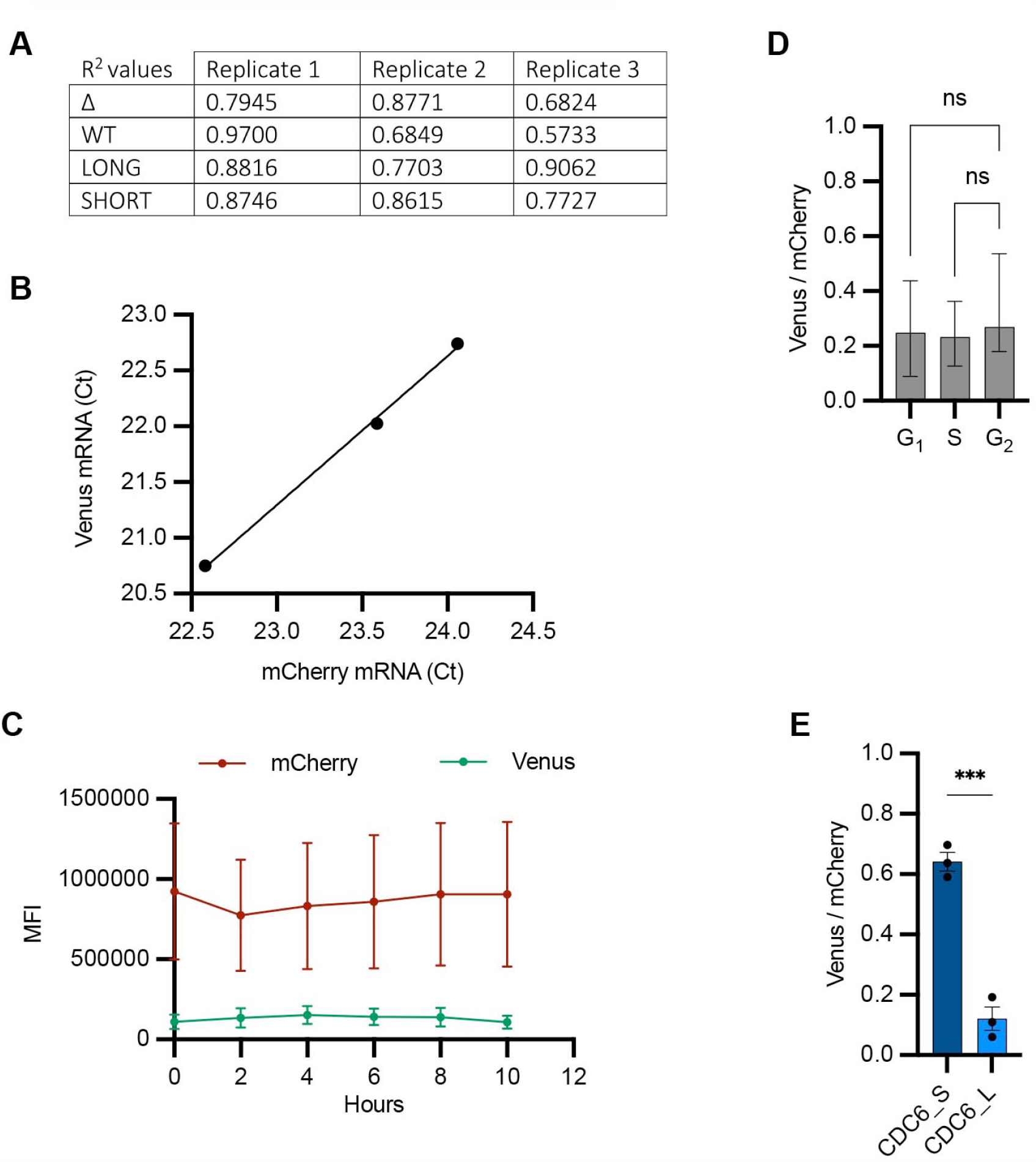
**(A)** R^2^ (coefficient of determination) values to indicate goodness of fit of a simple linear regression between mCherry and Venus single-cell MFI values. Data relative to **Figure 2B. (B)** RT-qPCR of mCherry and Venus mRNAs from three RNA extracts of U2OS cells transfected with Δ reporter. Ct values are mean between three technical replicates of the amplification reaction. **(C)** Quantification of mCherry and Venus MFI from U2OS cells transfected with the Δ reporter and imaged at the indicated time points. Mean and s.e.m. (n=8 cells) shown at each time point. **(D)** Single-cell Venus/mCherry MFI values from U2OS^CDK2^ cells transfected with Δ reporter grouped by intervals of CDK2 activity (**Figure 4A**, *right*) and plotted as median and 95% CI. n = 43 cells analysed. Kruskal-Wallis with Dunnett’s multiple comparisons test; ns, not significant. **(E)** Mean and s.e.m. of median Venus/mCherry MFI ratios from U2OS cells transfected with constructs containing long or short CDC6 3’UTR from three biological replicates. n ≥ 119 cells per condition. Unpaired t-test; ***, p=0.0005.

**Figure 4—figure supplement 1.**
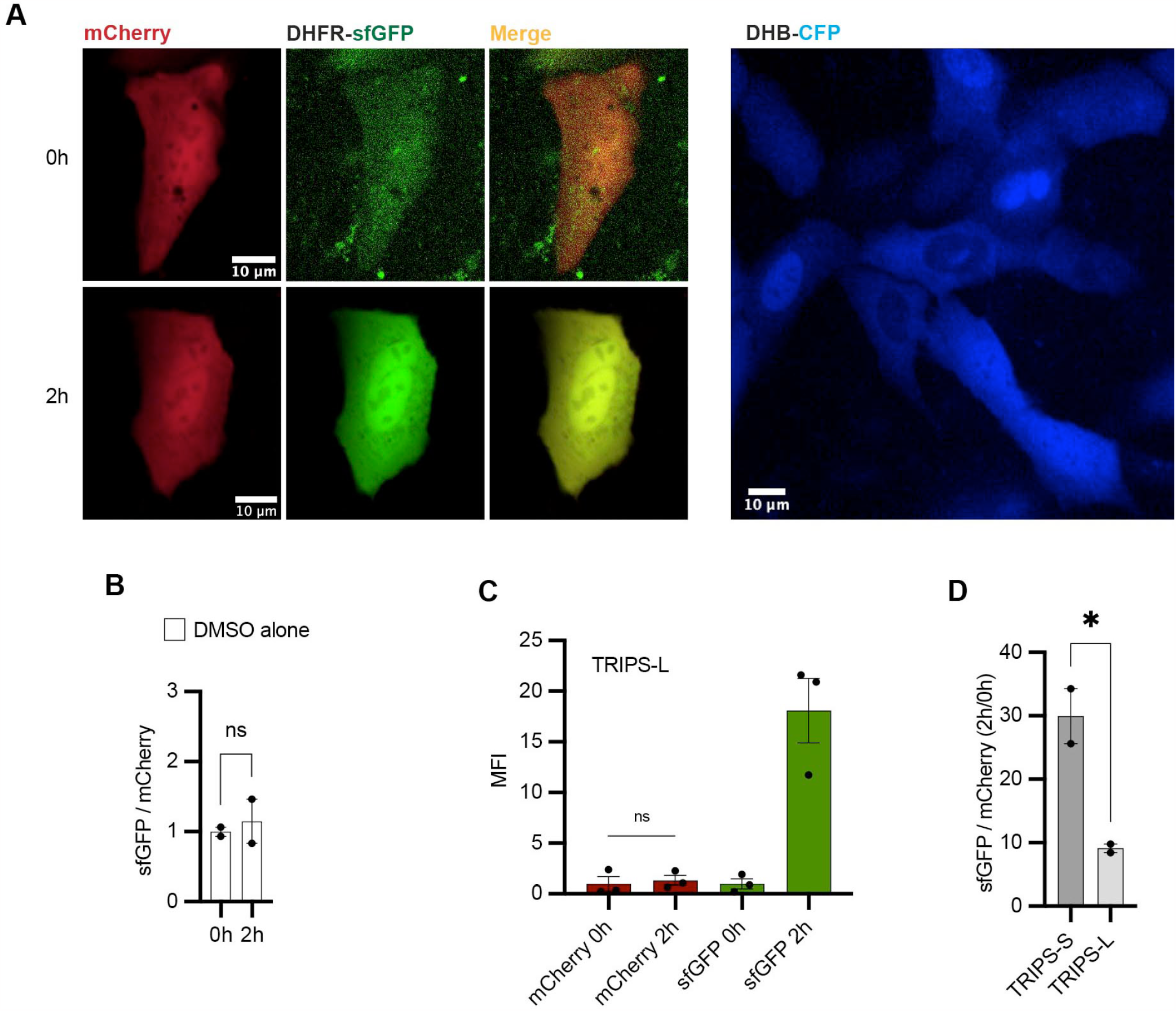
**(A)** *Left*. Representative U2OS cells transfected with TRIPS-S and imaged at 0h and 2h of 50μM TMP treatment. *Right*. Representative U2OS^CDK2^ cells. **(B)** TRIPS assay performed in U2OS cells transfected with TRIPS-Δ and treated with DMSO for 2h. Mean and s.e.m. of median sfGFP/mCherry ratios from two biological replicates, with baseline at 0h. n = 95 cells analysed. **(C)** Graph using data from **Figure 4D** showing MFI of mCherry and sfGFP at 0h and 2h of TMP treatment. Mean and s.e.m. of median MFI values from three biological replicates. **(D)** TRIPS assay performed in transfected U2OS cells and imaged at 0h and 2h of 50μM TMP treatment. Ratios between the median of single-cell sfGFP/mCherry ratios at 2h and that at 0h of TMP treatment were calculated for two biological replicates and are shown as mean and s.e.m.. n ≥ 343 cells analysed per condition. **(B), (C), (D)** Unpaired t-test. ns, not significant; *, p<0.05.

**Figure 6—figure supplement 1.**
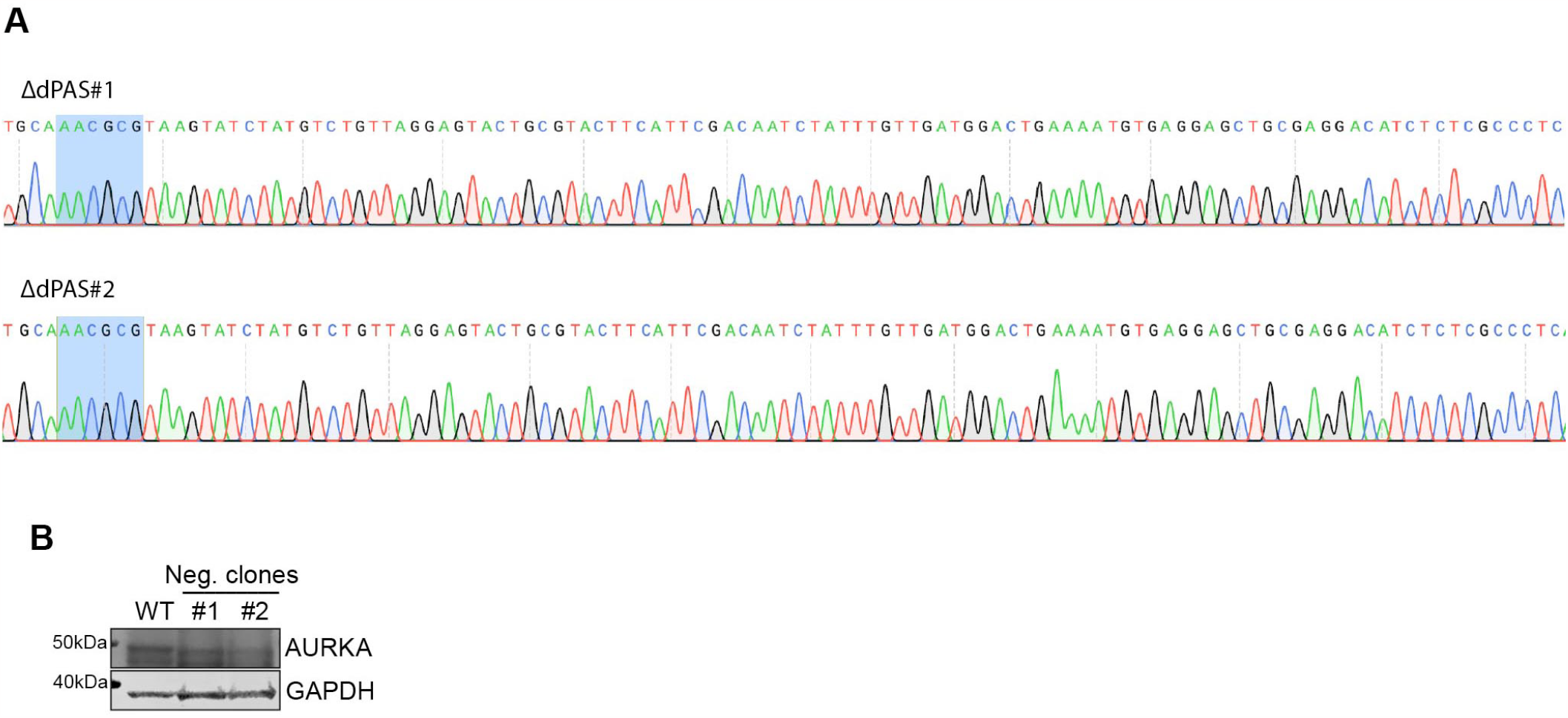
**(A)** Sanger sequencing validation of the mutated dPAS element in two cell lines. **(B)** Western blot showing similar AURKA protein expression in unsynchronized WT or U2OS cell lines that were not mutated following CRISPR editing (#1, #2).

## Notes

### Competing Interest Statement

The authors have declared no competing interest.

